# Servo-Actuated 3D-Printed Disposable Microvalves for Automated, Scalable Organoid Culture in Standard Incubators

**DOI:** 10.64898/2026.06.16.732526

**Authors:** Mojtaba Zeraatkar, Drew Ehrlich, Sebastian Hernandez, Hunter E. Schweiger, Mirella Pessoa de Melo, Eliot Wachtel, Demir Ozcakir, Spencer T. Seiler, Kateryna Voitiuk, Yohei Rosen, Colleen Josephson, Mohammed A. Mostajo-Radji, David Haussler, Sofie R. Salama, Mircea Teodorescu

## Abstract

Automation of organoid and cell culture processes is essential for achieving scalable and standardized experimentation in regenerative medicine and stem cell research. However, existing microfluidic platforms often rely on complex setups, limiting their integration within standard incubator environments. To address these challenges, we developed a compact, scalable multi-well platform featuring 3D-printed, servo-actuated disposable microvalves for fully automated media and drug exchange. This design eliminates the need for external pressure sources and control channels, providing a simplified and cost-effective solution for organoid culture. The platform integrates an internet-connected microscopy module with a motorized XYZ stage, allowing continuous, real-time imaging of individual wells directly within the incubator. It supports precise and reliable fluid handling under physiological conditions, improving throughput, reproducibility, and accessibility. We validate the platform through bench-top testing and in both mouse and human organoid models. Morphological analysis, immunohistochemistry (IHC), and qPCR demonstrate comparable viability, growth, and gene expression profiles between automated and manual culture conditions. These results establish a robust and scalable framework for fully automated organoid culture, offering a simplified and accessible alternative to conventional microfluidic systems with broad applications in regenerative medicine, drug discovery, and scalable biological screening.

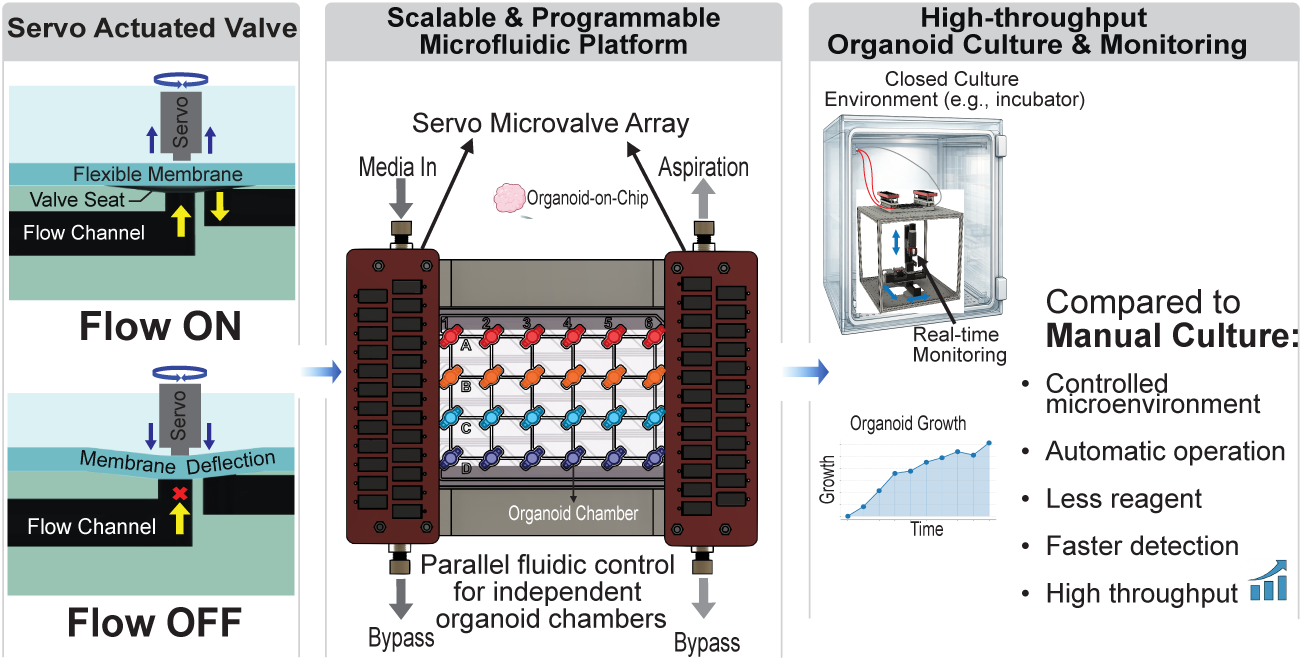

## 1. Introduction

Brain organoids hold significant promise for advancing our understanding of human brain development, modeling neurological disorders, enabling drug discovery, and supporting applications in regenerative medicine [34, 1]. However, traditional organoid culture methods suffer from limited biological fidelity, as they rely heavily on manual handling over months-long workflows and lack precise control over culture conditions. [8, 13]. The limitations reflect broader challenges in drug discovery, where studies estimate that the median research and development (R&D) cost to bring a single cancer drug to approval by the U.S. Food and Drug Administration (FDA) is approximately $648 million, with a median development time of 7.3 years and failure rates exceeding 96% [33, 48, 12]. These high failure rates are largely attributed to the inability of conventional in vitro and animal models to accurately recapitulate the complexity of human tissues.

The challenges highlight the need for advanced, automated, and scalable platforms capable of delivering precise and reproducible control over the cellular microenvironment. Robotic liquid handling platforms are commonly used as the gold standard for ultra-high-throughput cell culture and screening, offering precise and reproducible liquid dispensing across large sample sets [41, 24]. However, these platforms typically rely on discrete liquid handling steps and are not designed to provide continuous or spatially resolved modulation of the cellular environment, which is essential for organoid development and maturation [7, 50].

Organoid culture in microfluidic chips is emerging as a powerful approach to address these limitations through precise and dynamic control over the organoid microenvironment [29, 15]. Recent efforts have explored scalable organoid culture using microfluidic and perfusion-based platforms [17, 44, 39]. Perfusion bioreactors and microfluidic systems have demonstrated improved nutrient delivery and reduced variability compared to static culture. In addition, vascularized organoid-on-chip systems have enabled more physiologically relevant tissue models by integrating perfusable vascular networks within microfluidic environments [35, 3]. High-throughput organoid platforms based on microwell arrays and microfabricated devices have enabled parallelization, but often rely on passive diffusion or limited fluidic control [16, 38]. Despite these advances, many existing platforms still rely on external fluidic infrastructure and complex actuation systems, which can limit scalability and integration within standard incubator environments. Furthermore, most existing microfluidic organoid systems rely on reusable components that must be sterilized between experiments, introducing contamination risk and limiting experimental flexibility.

To enable programmable fluid control, microfluidic systems rely on integrated actuation components such as valves and pumps to regulate flow routing and timing. Various actuation strategies have been explored, including mechanical [32, 45], pneumatic [18], electrokinetic [25, 22], phase-change mechanisms [49], and external forces [5, 6]. Among these, pneumatic actuation remains the most widely adopted approach due to its reliability and compatibility with soft lithography-based fabrication [42]. However, pneumatic systems require additional control channels, tubing, external valves, and pressure sources. These requirements introduce significant complexity, cost, and system footprint, resulting in a bulky system that limits their suitability for scalable organoid culture inside standard incubators.

Fabrication methods further influence the scalability and accessibility of microfluidic systems. Traditional soft lithography techniques offer high precision but require cleanroom infrastructure and multi-step fabrication processes. In contrast, additive manufacturing techniques such as stereolithography (SLA) and digital light processing (DLP) enable rapid fabrication of complex three-dimensional geometries with reduced assembly requirements and increased design flexibility [2, 4]. While these methods facilitate rapid prototyping and system integration, incorporating active components such as valves and pumps into multiplexed architectures remains a significant challenge. Recent work has explored 3D-printed pneumatic valves as alternatives to PDMS-based systems [28, 37, 36], but these designs still inherit the complexity associated with pneumatic control.

Here, we present a compact and scalable automated organoid culture platform based on 3D-printed disposable microvalves actuated by miniaturized servomotors. The system integrates independently addressable fluidic control with longitudinal imaging and remote operation through an Internet-of-Things (IoT) enabled control framework. Compared with pneumatically actuated valve systems, this platform substantially reduces system complexity and footprint while supporting programmable, parallelized organoid culture within standard incubator environments.

## 2. Results and Discussion

### 2.1. Microvalve Design and Performance

Microvalves have been central to the widespread adoption of PDMS-based microfluidics, where the material’s intrinsic elasticity provides a natural basis for flow control.[42]. Consequently, significant efforts have focused on developing 3D-printed microfluidic systems with comparable valve capabilities to simplify fabrication and improve accessibility. The first 3D-printed microvalves were introduced by Nordin [36], and Folch [2], both based on pneumatic actuation. Such approaches, however, require high-resolution DLP printing and custom resin formulations, placing research-grade microfluidic printers (15–30K USD) out of reach for most laboratories [40]. Here, we adopt a fabrication strategy that enables 3D-printed microvalves to be produced using widely accessible printers and actuated via servo-controlled electromechanical mechanisms, as reported by Winkler et al. [47]. The resulting multilayer valve architecture builds upon conventional pneumatic designs while replacing pressure-driven actuation with electromechanical control, enabling scalable manufacturing and single-use operation.

An overview of the automated organoid culture platform is presented in Fig. 1, where the system operates inside a standard incubator while communicating through an IoT-based architecture [30] with external devices, including syringe pumps and microscopy modules.

**Figure 1:**
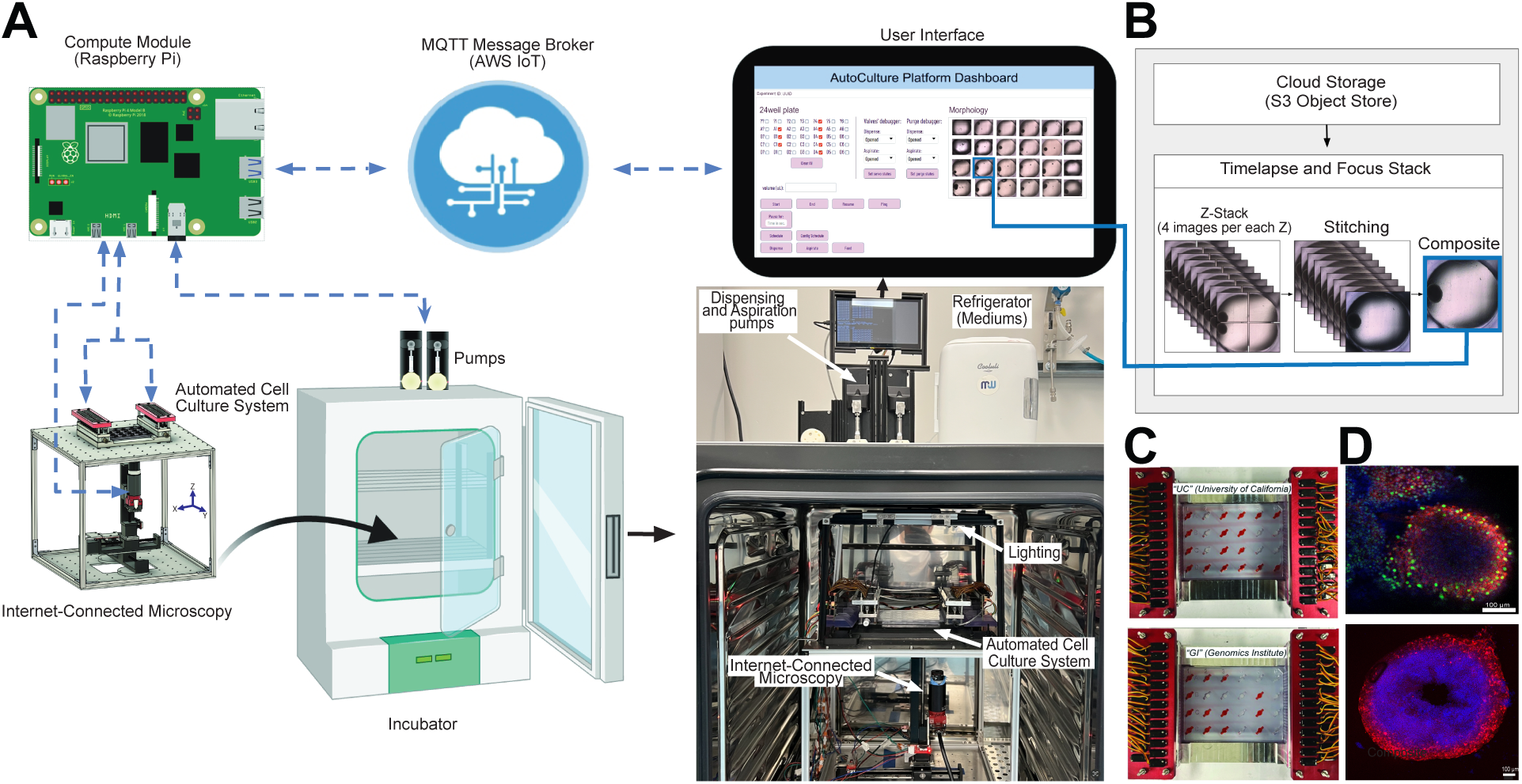
Automated Microfluidic Organoid Culture Platform. (**A**) Overview of the compact automated cell culture system integrated inside a standard incubator, including fluidic control, user interface, IoT, and internet-connected microscopy. **(B)** Remote monitoring and data workflow enabling live morphology assessment inside incubator. Z-stack images are captured at defined axial intervals, stitched across multiple regions of each well, and combined into composite focus stacks, which are further processed into long-term time-lapse datasets for continuous growth analysis (see Supplementary Video S2). **(C)** Images showing “Dry run” test results of the developed system with a colorant dye; the system writes predefined patterns (“UC” for University of California and “GI” for Genomics Institute) during automated operation (see Supplementary Video S1). The developed system is programmable and provides exchanging medium and drugs from a single well with different volumes and flow rates. **(D)** Immunohistochemistry (IHC) analysis of organoids following completion of the experiment, confirming tissue identity and structural integrity.

Conventional microfluidic systems are primarily based on pneumatic actuation for valve operation (Fig. 2A), requiring additional control channels, tubing, fittings, external valves, and pressure or vacuum sources, which increase system complexity and footprint. In contrast, our platform employs electromechanical actuation using low-cost miniature servomotors (FH-1083, Flash Hobby), each costing $3.7. The system integrates 50 valves to enable media and drug exchange across a 24-well plate: 24 valves for media delivery, 24 for aspiration of spent media, and two purge valves. The platform is organized into six linked modules: (1) a 24-well plate; (2) a microvalve system; (3) servomotors and their holders; (4) a structural fixture; (5) a control system; and (6) external components, including syringe pumps and a refrigerator.

**Figure 2:**
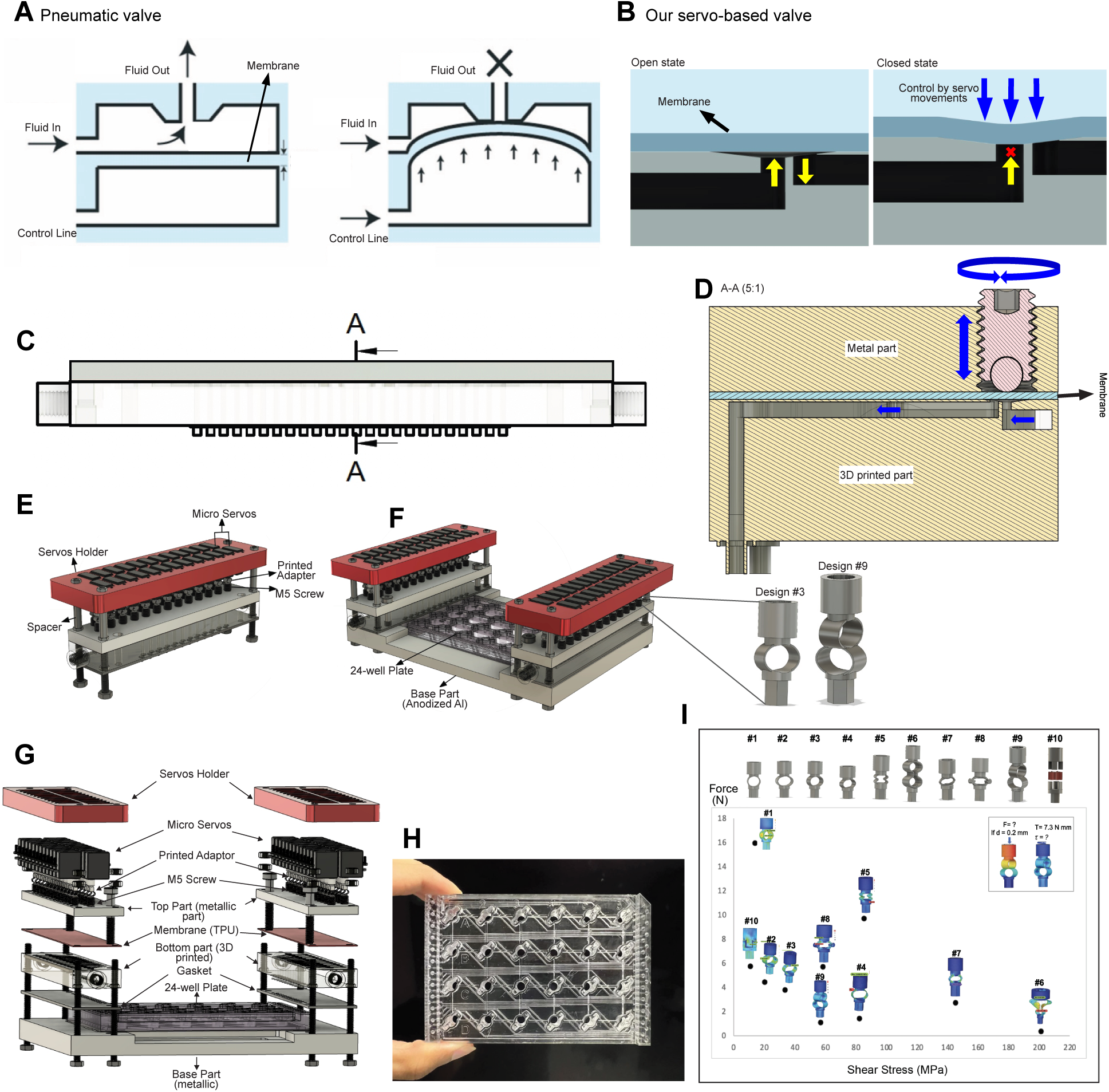
System Overview. (**A**) Schematic diagram of a pneumatic valve; two sets of fluidic and control (air pressure) channels are needed. **(B)** Schematic diagram of our servo-based microvalve system with cut-away schematics that illustrate a membrane valve in open and closed states depending on movements of servo. **(C)** Front View of the valve system: the system consists of three main parts: Bottom part which contains fluidic channels and valve seats; Membrane: Flexible thermoplastic; Top part made of stainless steel. **(D)** Cross section showing the mechanism of a microvalve. Servos positioned on top of the system and linked to the top part using a ball tip screw and a flexible adaptor (I) which convert the rotational movements of servo into a linear movement. Closing and opening of each valve carry out using a rotation angle of 90 degree CW and CCW, respectively. **(E)** Schematic showing our microvalves system. **(F)** Schematic showing assembled automated cell culture system; The design including a 24-well plate chip assembled between microvalve system and the base part using gaskets in order to prevent fluid leaking. Exchanging media is facilitated through the utilization of microservos (50 valves) positioned on both sides of the 24-well plate chip. **(G)** An exploded view of the system showing all parts. **(H)** 24-well plate chip, and **(I)** FEA analysis for choosing the optimum design for M5 adaptor part. A variety of different designs considered in order to find the optimum design based on minimum force that need for 200 µm displacement and minimum shear stress under a torque of 7.3 N mm applied by a servo.

The valve design consists of a multilayer architecture comprising three main components (Fig. 2B–D). The bottom layer includes fluidic channels and valve seats with a diameter of 4 mm and a depth of 200 µm. A flexible TPU membrane with a thickness of 600 µm serves as the intermediate layer and deflects into the valve seat (Fig. 2B) through a coupling mechanism integrated into the metallic upper layer of the microvalve. The coupling system includes M5 ball-tip screws (M5-0.8 × 8 mm) and M5 adapters. The detailed fabrication process is described in Supplementary Fig. S1. Valve actuation is achieved through 90° clockwise and counterclockwise rotation of the servomotor, which is converted into linear displacement via the coupling mechanism, enabling opening and closing of the valve.

The printed adapters play a critical role in actuation and is designed to provide flexibility in the z-axis to minimize permanent loading on the servomotor, while maintaining rigidity in the xy-plane to withstand shear stresses induced by servo torque. A total of ten candidate geometries were evaluated based on mechanical performance and manufacturability. Designs 9 and 3 successfully passed the mechanical stress testing (Fig. 2I), demonstrating reliable performance under ambient and high-humidity incubator conditions, respectively (see section 2.3). Finite element analysis (FEA) results are provided in Supplementary Fig. S2.

A 24-well plate chip was designed to support 3D organoid culture, with a footprint of 85.1 mm × 127.4 mm. The layout consists of four rows (A–D) and six columns (1–6) (Fig. 2H). Each well is individually addressable through square channels with dedicated inlets, outlets, and an organoid chamber. The effect of well geometry in this platform was previously characterized using Computational Fluid Dynamics (CFD), demonstrating transient, low-velocity flow regimes that minimize shear stress on organoids [39]. Media flow in each well is independently controlled using integrated pumps and microvalves, enabling precise and reproducible exchange conditions. The plate integrates into the overall system as described in section 4.5 and shown in Fig. 2.F, with microvalves positioned on both sides. The design enables straightforward disposal of both the valves and the 24-well plate after each experiment. It further facilitates seamless integration with laboratory instruments, including an in-incubator imaging system (Fig. 1), through an Internet of Things (IoT) framework.

### 2.2. System Control and Valve Characterization

The automated cell culture system is controlled by a microcontroller (Raspberry Pi 4), which coordinates all subsystems, including fluidic components (valves and pumps), electromechanical actuators, custom PCBs, and the imaging system. Communication between devices is managed through an Internet of Things (IoT) architecture in which cloud-based MQTT brokers facilitate data exchange, storage, and processing, as described in [30, 43]. All subsystems operate through custom-developed software [27] that links users, devices, and data, providing experiment scheduling, device control, real-time monitoring, and inter-device communication between fluidic systems, imaging hardware, and cloud services. Through the graphical user interface (Fig. 1A), the well plate is displayed as a schematic 6 × 4 grid (wells A1–D6) that mirrors the physical layout. The user selects one or more target wells directly from this grid and, for each selected well, independently specifies the action (dispense, aspirate, or feed) and its parameters, including the volume and timing. The complete per-well schedule is then translated into device-level commands that are distributed to the corresponding microcontrollers and executed autonomously. The same interface also streams per-well morphology images for continuous visual monitoring of long-term experiments.

A schematic of the control architecture is shown in Fig. 3A. Upon receiving dispense, aspirate, or feed commands from the user via the user interface, the system selectively actuates the appropriate microvalve to open/close a fluidic path to the target well. Valve actuation is achieved via pulse-width modulation (PWM) signals generated by the servo control PCB, which drive the servomotor to rotate 90° counterclockwise (CCW), thereby opening the corresponding valve. For media delivery, once the fluidic path is established, the syringe pump draws the programmed volume of media from the reservoir stored in a refrigerator and dispenses it at a specified flow rate through the open fluidic path into the selected well. After delivery, the valve is closed by rotating the servomotor 90° clockwise (CW), isolating the well before proceeding to the next command. For aspiration, the corresponding valve is similarly opened via a 90° CCW rotation, and a second syringe pump removes spent media from the well by aspiration through the open fluidic path. The aspiration height is controlled to a predefined threshold (5.9 mm from the bottom of the well) to ensure sufficient residual media remains, thereby preventing desiccation of the well.

**Figure 3:**
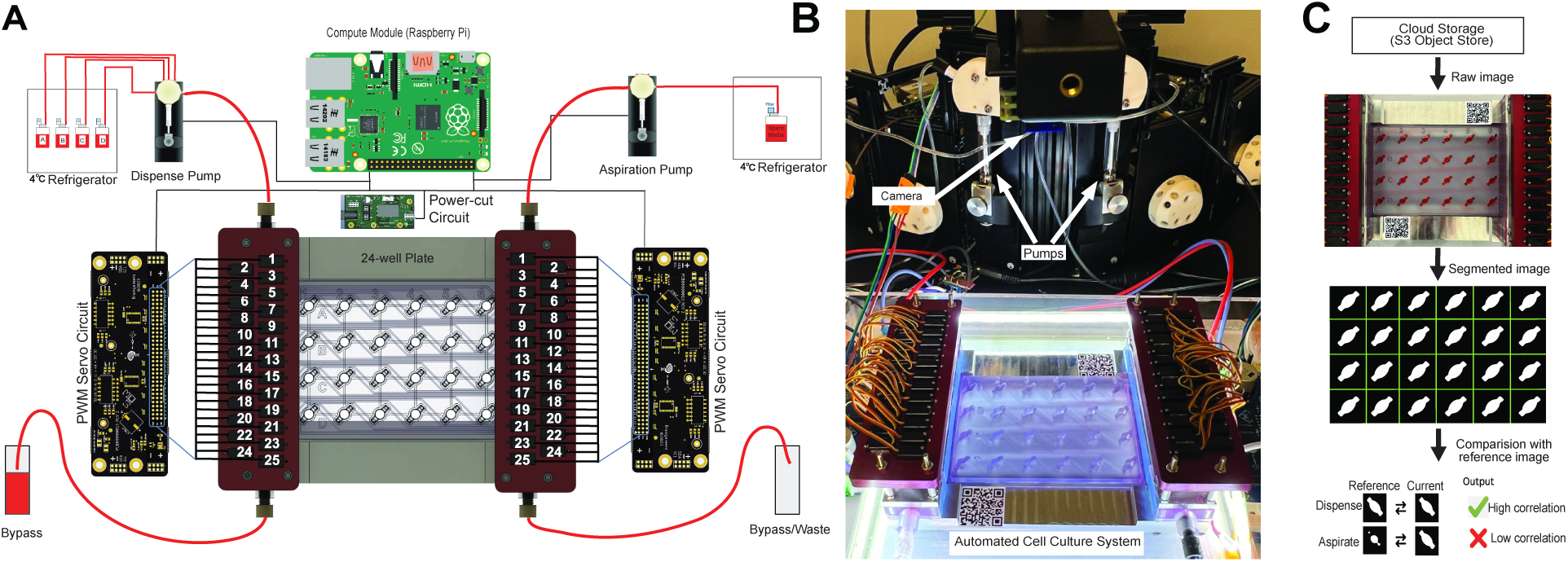
Control architecture and dry-run testing of the automated cell culture system. (**A**) Schematic of the control part of automated cell culture system. **(B)** Bench-top stress testing and dry-run validation using red dye to evaluate aspiration and dispensing reliability. The camera was positioned perpendicular to the well plate and acquired images after each aspiration and dispensing command for every well. **(C)** Algorithmic workflow used for reliability assessment based on images captured after each feeding cycle and correlation analysis, with quantitative results summarized in Supplementary Table S1.

The system incorporates two dedicated purge valves, which serve dual functions: (i) flushing residual media or reagents from fluidic lines during switching between different media, and (ii) acting as a pressure relief mechanism to prevent pressure buildup during operation cycles. Additionally, a custom power-switching circuit disconnects power to the servomotors when the system is idle, thus minimizing permanent mechanical loading on the servo shafts during long-term experimentation.

### 2.3. Bench-top Validation and Long-term Reliability

The valve architecture, including miniature servomotors, a coupling mechanism, a flexible membrane, and a multilayer, multi-material structure, requires robust long-term stability and reliability for cell culture applications. After system assembly (section 4.5), the platform was subjected to a series of mechanical stress and bench-top validation tests. First, the system was tested for stress, during which all 50 valves were repeatedly opened and closed every 5 s to evaluate mechanical performance. The platform completed more than 12,000 actuation cycles per valve without mechanical failure, demonstrating long-term operational reliability of the valve design and actuation mechanism under repeated stress conditions. Given a typical feeding frequency of 4 times daily, the system requires approximately 240 actuations per valve per month.

To evaluate whether the automated platform can reliably support long-term media exchange, the performance of the system was assessed through bench-top experiments in which a colorant red dye in a 24-well plate was automatically exchanged every hour over a period of 4 days. In each cycle, 150 *µ*L of dye was aspirated from each well and replaced with 150 *µ*L of fresh dye. A camera positioned above the plate captured images after each individual feed and aspiration event (Fig. 3B). For each of the 24 wells, a sequence of *N* aspiration and dispensing commands was executed. Using the computer-vision pipeline described in Section 4.7 and illustrated in Fig. 3C, each operation was classified as successful or failed, enabling calculation of per-well and aggregate success rates. The results per well are summarized in Supplementary Table S1.

Dry-run experiments across all 24 wells demonstrated high reliability of automated media handling, with a dispensing success rate of 99.9% (CV = 0.35%) and an aspiration success rate of 98.1% (CV = 2.9%). The combined operational success rate was 99%, confirming robust and reproducible system performance suitable for long-term automated cell culture. These success rates specifically reflect the performance of the valves in directing flow to and from the target wells. The precision of volumetric delivery is determined by the syringe pumps (CV ≤ 0.05%). Aspiration failures were observed in a small subset of valves (see Supplementary Table S1), attributed to a temporary adhesion of the flexible membrane to the valve seat. Unlike dispensing, where positive pressure helps lift the membrane from the valve seat, aspiration relies on suction, which can block flow if the membrane adheres. To address this, a brief positive pressure pulse was introduced prior to each aspiration command, facilitating membrane release and enabling reliable spent media removal.

### 2.4. Automated Organoid Culture Across Mouse and Human Models

#### 2.4.1. Mouse organoids

The automated cell culture platform was evaluated using mouse embryonic stem cell (mESC)-derived organoids to assess its ability to support long-term maintenance and development under fully automated conditions. Organoids were generated following the protocol described in Section 4.8. As an initial validation step prior to full-plate biological experiment, mouse dorsal organoids in day 29 were first cultured in a limited subset of wells (four wells) to evaluate system compatibility and media exchange performance. Organoids were cultured for 7 days (D29–D36). The system executed a fully programmed feeding protocol in which each well was initially fed with 250 *µ*L of media, followed by automated media exchange every 12 hours. During each cycle, the spent media was aspirated and replaced with 150 *µ*L of fresh media. In parallel, control organoids were maintained under manual feeding conditions. An integrated in-incubator imaging system, equipped with a motorized XYZ stage and LED illumination, enabled high-resolution brightfield imaging across multiple Z-planes at hourly intervals. Morphological observations (Fig. 4B) and IHC analysis (Fig. 4E) confirmed maintenance of organoids with morphology and marker expression comparable to static controls. At this stage, organoid growth was limited, as previously reported in [14].

**Figure 4:**
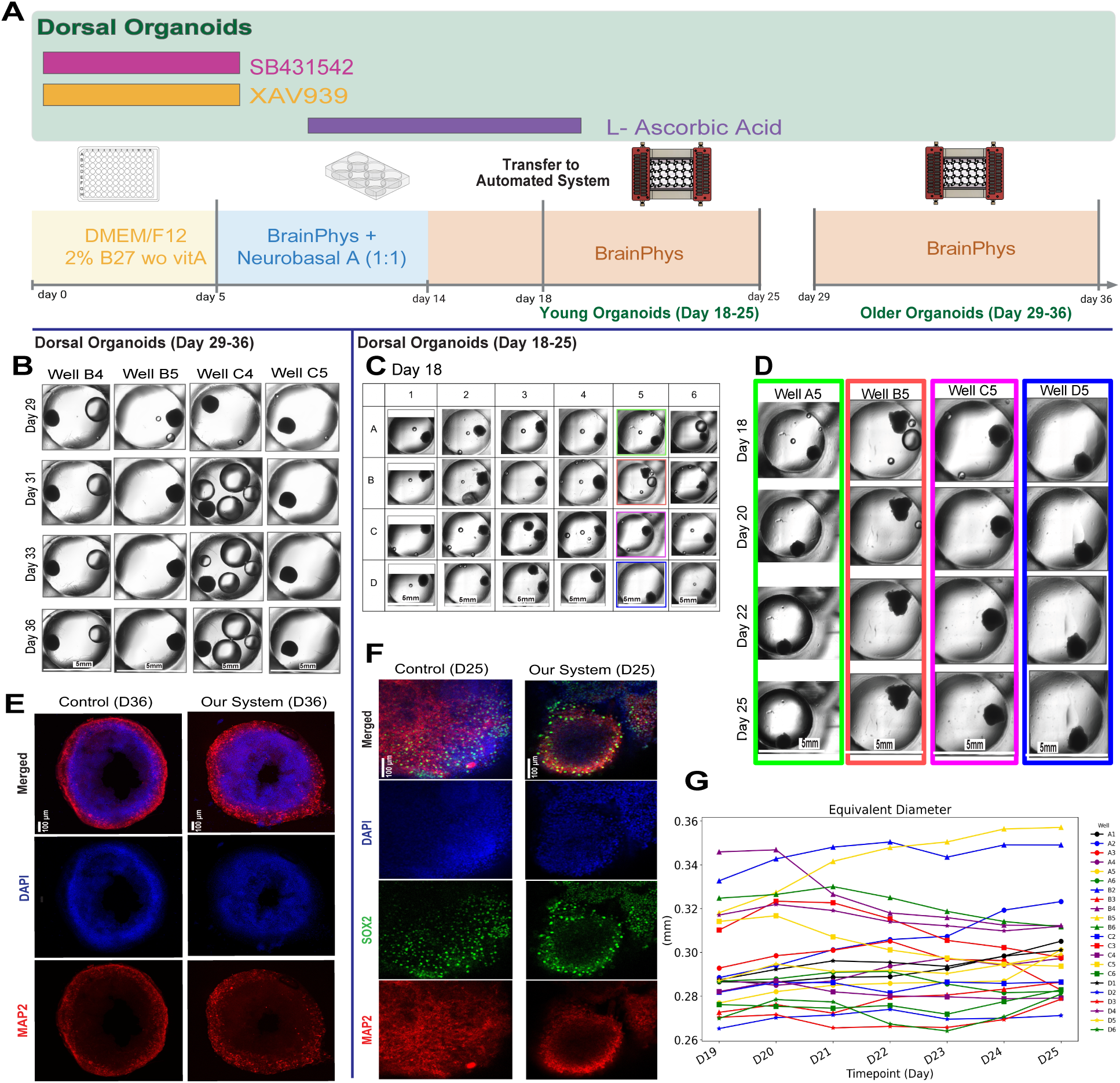
Experimental workflow, morphological assessment, and immunohistochemical (IHC) characterization of dorsal mouse organoids cultured in automated cell culture system compared with static control organoids. (**A**) Timeline of dorsal mouse organoid experiments, including plating, media changes, and endpoint collection. Two experiments using different ages of organoids were performed and compared with static controls. **(B)** Brightfield images showing organoid morphology (Day 29-36) across four wells during automated culture with IHC analysis in **(E)** compared with static control. **(C)** Brightfield images showing organoid morphology in day 18 across all 24 wells, with a representative column in **(D)** illustrating morphological progression over the culture period (Day 18–25), including growth analysis in **(G)**. Wells B1 and C1 and, the second organoids in wells C3 and D3 were excluded from statistical analysis because of partial organoid capture. **(F)** IHC analysis comparing organoids cultured in the automated platform versus static controls at Day 25. Staining for neural markers (MAP2, SOX2, DAPI) confirms preservation of structural integrity and expected cell-type composition.

Following the initial evaluation, the system was expanded to larger well counts by replacing the disposable 24-well plate and microvalve components. The use of disposable components enabled rapid system reconfiguration between experiments, with fabrication costs of approximately $20 per 24-well plate and $8 per microvalve fluidic unit. Critically, each new experiment begins with single-use sterile components, eliminating the cross-contamination risk inherent to reusable microfluidic systems, which is a particular advantage when culturing distinct organoid lines in parallel. This disposability also eliminated cleaning and sterilization downtime between experiments, allowing immediate system redeployment after each run.

For the second independent mouse organoid experiment, younger organoids (D18) were transferred into the automated culture system with one organoid per well; wells C3 and D3 were unintentionally plated with two organoids each. Organoids were then cultured for 7 days (D18–D25) using the automated feeding protocol described in the initial experiment.

Fig. 4C shows representative morphologies for all wells in day 18, with a column highlighted in Fig. 4D over the 7-day timeline. Statistical analysis of organoid growth is presented in Fig. 4G, and time-lapse imaging of two representative wells in Supplementary Video S3 further demonstrates organoid growth and morphological stability throughout the culture period. To evaluate biological performance, immunohistochemistry (IHC) analysis was performed following harvesting at day 25. As shown in Fig. 4F, staining for neural lineage markers (MAP2, SOX2) and nuclear marker (DAPI) demonstrated that organoids cultured in the automated system maintained structural integrity and expected cellular organization, comparable to manually maintained controls. These findings are consistent with previously reported organoid differentiation profiles [14].

#### 2.4.2. Human organoids

To demonstrate the applicability of the automated platform to more complex biological systems, human embryonic stem cell (hESC)-derived organoids were cultured under fully automated conditions. Organoids were generated following the protocol described in Section 4.8 and transferred into a 24-well plate on day 16. A 9-day experiment was conducted to evaluate the system’s ability to deliver distinct media formulations and execute distinct feeding protocols in parallel across the plate.

Organoids were distributed across a 24-well plate, with each row assigned to a specific feeding protocol. Rows A and B were subjected to a lower-frequency feeding regimen (media exchange every 12 hours), while rows C and D received higher-frequency feeding (every 6 hours). Each well was initially filled with 200 *µ*L of AggreWell EB formation medium. Automated media exchange was performed by aspirating and dispensing 100 *µ*L per cycle.

From day 2, the system switched to DMEM/F12 + Glutamax + N2 medium with the same feeding schedules. The internet-connected microscopy system acquired brightfield images of each well at defined z-stacks every 2 hours(Fig. 5B). Organoids across the entire plate exhibited sustained growth and expected morphological features over the 9-day culture period, with a representative column highlighted in Fig. 5C (see Supplementary Video S2 for a time-lapse). Quantitative analysis of organoid growth (Fig. 5D–G) showed a general increase in size across all conditions. While both feeding regimens supported growth, organoids cultured under higher-frequency feeding exhibited increased growth variability across wells, suggesting greater sensitivity to more frequent media exchange. Gene expression analysis (Fig. 5H) was performed using qPCR and normalized to control H9 organoids. Notably, the control samples were approximately 10 days more mature than the organoids cultured in the automated system. Consequently, reduced expression of FOXG1 and increased expression of PAX6 were observed in automated samples, consistent with a more progenitor-like developmental state characteristic of earlier-stage organoids. This trend aligns with established developmental trajectories in human cortical organoids, where early-stage cultures are enriched in PAX6+ neural progenitors and progressively transition toward FOXG1-associated forebrain identity during maturation [31]. Importantly, no substantial differences in gene expression profiles were observed between low and high frequency feeding conditions, indicating that both protocols support similar lineage specification outcomes.

**Figure 5:**
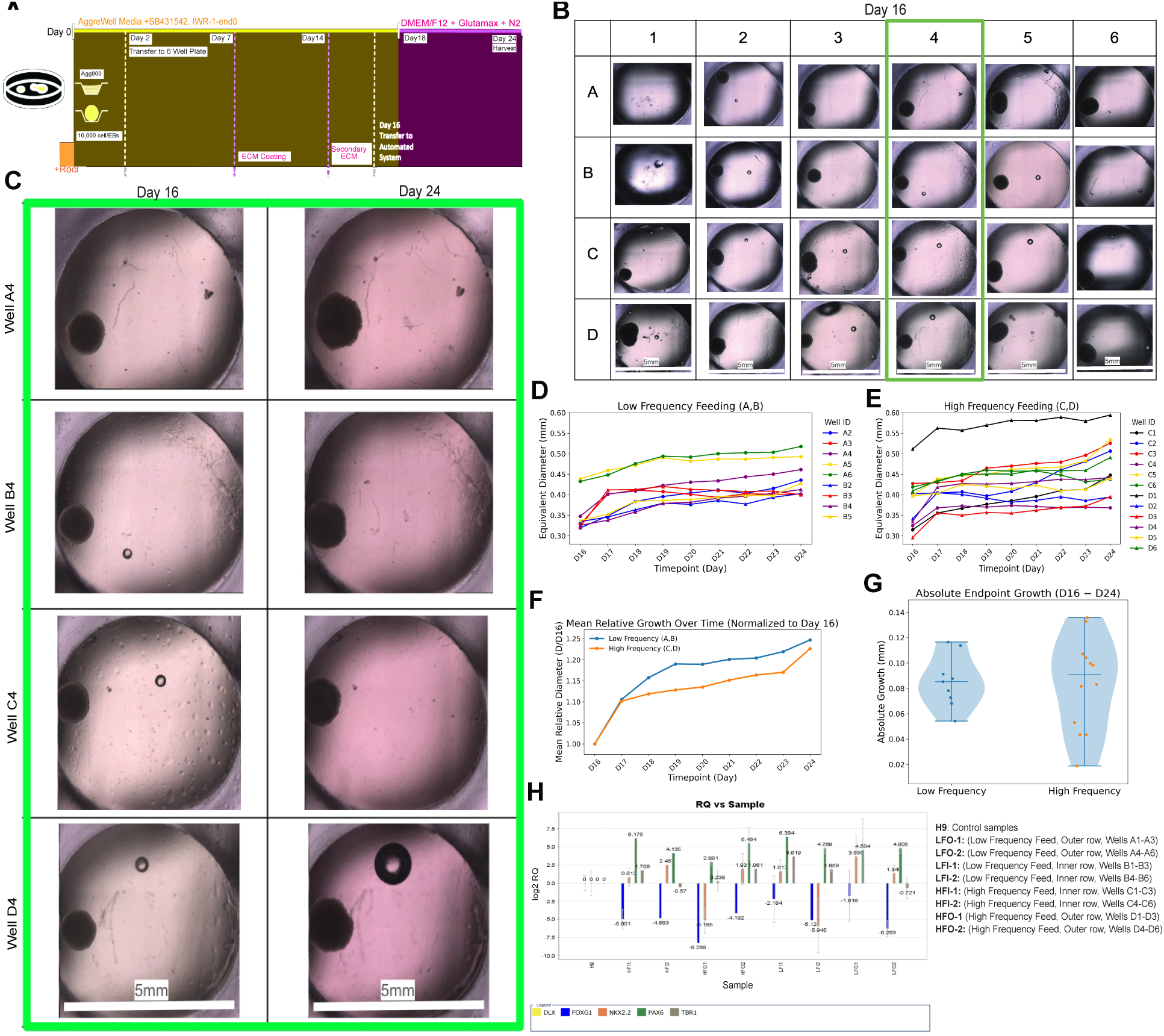
Experimental design, organoid morphology, growth dynamics, and gene expression analysis under different feeding protocols. (**A**) Timeline of the 9-day human organoid experiment, including plating, different medium, and endpoint collection.**(B)** Brightfield images showing organoid morphologies across all 24 well in day 16, with a representative column in C. **(D–E)** Growth trajectories of individual wells under low-frequency feeding (D) and high-frequency feeding (E), showing temporal changes in organoid size. **(F)** Mean organoid growth normalized to D16 for both low and high frequency feedings. **(G)** Comparing low-frequency and high-frequency feeding conditions. **(H)** Relative gene expression of selected markers (DLX, FOXG1, NKX2.2, PAX6, and TBX1) across experimental conditions, including control samples and, low and high frequency feeding groups from inner and outer well positions.

Osmolarity measurements (Supplementary Table S2) showed higher evaporation rates, particularly in wells located in row A. This edge effect is a well-known phenomenon in multi-well plate cultures, where evaporation gradients are typically higher at the corners and edges of well plates [26, 46]. Notably, higher-frequency feeding mitigated these effects, particularly in row D, by more effectively compensating for evaporative losses. These findings highlight an additional advantage of our automated cell culture platform in maintaining more uniform culture conditions across the plate and reducing spatial variability. This adaptive capacity to detect and mitigate spatial evaporation gradients through programmatic feeding frequency adjustment represents a capability that is difficult to achieve with manual workflows, where feeding intervals are typically fixed and spatially uniform regardless of well position.

## 3. Conclusion

Automation is a key driver for achieving standardized and reproducible experimental conditions in organoid research. In this work, we present an automated microfluidic platform that enables precise and scalable culture of mouse and human organoids. Our results demonstrate that the system can reliably support long-term organoid maintenance across multiple biological models, and its potential for reproducible and scalable in vitro systems.

By integrating 3D-printed disposable microvalves with programmable control, the platform enables robust, reproducible, and independently addressable media exchange for each well while maintaining biological performance comparable to conventional manual culture. The use of single-use sterile components eliminates cross-contamination risk between experiments and removes sterilization downtime, enabling rapid reconfiguration across independent experimental runs. In contrast to traditional robotic liquid handling systems, which represent the current gold standard, microfluidic valve-based platforms provide enhanced spatiotemporal control over fluid delivery within a compact and scalable architecture. The system further enables parallelization of experimental conditions within a multi-well format, allowing simultaneous evaluation of key variables, including feeding frequency, culture conditions, and intrinsic organoid-to-organoid variability within a single experiment. This capability is particularly important for organoid systems, where biological heterogeneity necessitates controlled and parallel experimentation.

The scalability of the automated microfluidic system, combined with its low fabrication cost and compatibility with standard laboratory infrastructure, positions this approach as a broadly accessible direction for high-throughput organoid culture and screening. The platform’s reliance on widely accessible desktop printers and low-cost components lowers the barrier to adoption for institutions with limited resources, broadening access to standardized and reproducible organoid experimentation. Future developments may integrate artificial intelligence and closed-loop feedback control, enabling dynamic adjustment of culture conditions based on real-time, organoid-specific responses. Such adaptive and individualized control of organoid development represents an important step toward more precise and reproducible models, with significant implications for regenerative medicine and stem cell research.

## 4. Materials and Methods

### 4.1. Design and Fabrication Process

All parts were designed using Fusion 360 (Autodesk Inc., San Francisco, CA, USA) and exported as STL files. The valve system components, including the bottom part (Fig. 2G) and the adapters used in the coupling system (Fig. 2I), were fabricated using Formlabs 3B+ printer and BioMed clear resin (Formlabs, Somerville, MA, USA). The 24-well plate (Fig. 2H) was fabricated using Asiga Ultra 3D printer (Asiga, Sydney, Australia) and Detax FREEPRINT Ortho resin (Detax GmbH, Ettlingen, Germany). The flexible membrane (Fig. 2G) was fabricated using an Ultimaker S5 3D printer (Ultimaker, Utrecht, Netherlands) with thermoplastic polyurethane (TPU) filament (Ultimaker, Utrecht, Netherlands). A 0.4 mm nozzle was used for printing, and the resulting TPU membrane had a thickness of 0.6 mm. The following printing parameters were applied: initial layer height of 200 µm, infill extrusion angle of 45° (which provided better membrane elasticity compared to 0° or 90°), 100% infill density, printing temperature of 223 °C, and a build platform temperature of 60 °C throughout the print. The print speed was 20 mm/s. The upper part of the valve design (Fig. 2G) and the fixture (base part, Fig. 2G) were CNC-machined from 316 stainless steel to minimize mechanical wear and improve corrosion resistance under the high-humidity conditions of the cell culture incubator (Billet Metal Craft, Santa Cruz, CA, USA). The servo holders (Fig. 2E) were CNC-machined from aluminum 6061 and anodized to prevent corrosion (Billet Metal Craft, Santa Cruz, CA, USA). The step-by-step fabrication process of the multilayer microfluidic platform is shown in Supplementary Figure S1.

Finite element analysis (FEA) was performed using Fusion 360 to evaluate the mechanical performance of the 3D-printed adapter (Fig. 2I), that couples the servo shaft to the M5 screw, using Autodesk Fusion 360. The analysis was done using the maximum torque generated by the servo motor (7.3 N·mm) and the required displacement to achieve membrane deflection (200 µm). All possible designs were analyzed assuming a consistent material model corresponding to BioMed clear resin, with mechanical properties defined based on reported material specifications [10]. The distributions of bending stress and shear stress were analyzed for multiple adapter geometries. The numerical results identified two candidate designs that minimized mechanical load on the servo shaft, thereby reducing the risk of long-term failure. The corresponding numerical simulation results are provided in Supplementary Figure S2.

These designs were further validated through benchtop testing and inside humid environmental conditions such as standard incubators. At room temperature, Design 9 demonstrated better performance by minimizing the load on the servo shaft. In contrast, under high-humidity conditions at 37 °C, Design 3 exhibited improved performance and successfully passed the mechanical tests.

### 4.2. Custom circuit boards (PCBs)

Custom servo control boards were designed to enable modular and scalable actuation. Each board is based on a pair of PCA9685 I^2^C-controlled PWM driver integrated circuits, enabling control of up to 32 servos per board. By utilizing configurable I^2^C addresses, up to 31 boards can be daisy-chained on a single bus, allowing control of up to 992 servos. The boards incorporate reverse polarity protection and standard 2.54 mm header pin arrays for power, data, and servo connections. To ensure reliable operation in high-humidity incubator environments, the boards were conformally coated with Super Corona Dope (MG Chemicals), and silicone gaskets were used between header pins to reduce moisture ingress.The servo control boards were mounted vertically on two fixtures in both sides of the system (Fig. 1A) to prevent droplet formation on the boards. The design separates logic and power domains, with independent voltage supplies and shared grounding, allowing selective power delivery to servos. This enables power gating to reduce continuous static load and heat generation when the system is idle.

To facilitate system integration, the boards feature symmetric power and I^2^C routing with standardized connectors, enabling straightforward daisy-chaining into a unified control bus. The PCBs were fabricated by Sierra Circuits (Sierra Circuits, Inc., Sunnyvale, CA, USA). Additional details of the servo and power control boards are provided in Supplementary Figure S3 and S4.

### 4.3. Biocompatibility and Cell Viability

BioMed Clear resin is an FDA-registered material, compliant with ISO 10993 for biocompatibility and USP Class VI standards, making it suitable for cell culture applications [11, 20].The BioMed Clear resin was used to fabricate the fluidic part (bottom part) and flexible adapters in the microvalve system.

To evaluate the biocompatibility of Detax FREEPRINT Ortho resin used for 24-well plate, we fabricated three one-well chips with different bottom surfaces: (1) printed resin bottom, (2) 3M^TM^ Microfluidic adhesive tape (9795R, 3M), and (3) glass substrate. The glass substrate fixed on the build plate and the part printed directly on glass. All parts were post-processed including washing in IPA and followed by Otoflash curint unit (Otoflash G171, NK Optik, Baierbrunn, Germany). The chips were then autoclaved and prepared for biological testing. Biocompatibility and cell viability were assessed using mouse organoids cultured for 11 days, with two organoids per well. As summarized in Supplementary Figure S6, organoids exhibited comparable viability and morphology across all substrates, indicating that FREEPRINT Ortho resin is biocompatible for cell culture applications. Among the tested substrates, adhesive tape and glass provided greater optical transparency, improving imaging quality for microscopy.

For the membrane, thermoplastic polyurethane (TPU) was selected due to its flexibility, mechanical durability, and established biocompatibility in biomedical environments [21]. TPU was used as the deformable membrane layer in the servo-actuated microvalve system.

### 4.4. Sterilization

The majority of components used in the automated cell culture system, including 3D-printed resin parts, were compatible with autoclave sterilization. Prior to assembly, all fabricated components were sterilized by autoclaving.

The only component of the microvalve system that was not autoclavable was the thermoplastic polyurethane (TPU) membrane. For these parts, which were integrated into the fluidic part (bottom part) of the valves, sterilization was performed using 10% hydrogen peroxide. Following system assembly (see section 4.5) and connecting tubings to syringe pumps, the fluid pathways were flushed with Spor-Klenz RTU sterilant (STERIS Life Sciences) to ensure sterility prior to biological experiments. After completion of each experiment, the valves and well plate were discarded and replaced with new sterile ones for subsequent runs.

### 4.5. Platform Assembly

The automated cell culture platform was assembled inside a biosafety cabinet. All components were sterilized prior to assembly. A step-by-step guide to the assembly process is provided in Supplementary Figure S5.

To provide precise volumetric flow and enable aspiration of spent media, syringe pumps (Tecan Cavro Centris, 1.0 mL glass syringe) were used as described in [39]. The syringe pumps, together with distribution valves, enabled delivery of different mediums to the 24-well plate via a single inlet line routed into the incubator, as well as aspiration from the wells through a dedicated outlet line for spent media. A compute module (Raspberry Pi 4) coordinated communication between the fluidic system and the microscopy setup within an IoT-enabled network. A touchscreen display was used to configure, edit, and execute experimental protocols.

### 4.6. In-incubator Microscopy

An internet-connected microscope was used to perform live imaging of the organoids. The microscope is a variant of the microscope described by Ehrlich et al. [9], designed with a three axis scanning stage to allow widefield live cell imaging of many wells in one experiment. In this case, the goal was to observe organoid cell culture across 24 separate wells over the course of multiple days. A high resolution USB camera (Allied Vision Alvium 1800U-500c) was used for these experiments. This camera has a full color ON Semi AR0521 CMOS sensor that can capture images that are 2592 x 1944 pixels in size.

The camera was mounted at the bottom of a 50mm lens barrel that has a 4X 0.1NA objective on the other end. The camera was connected to a computer via a USB cable and was controlled using the vmbpy Python library provided by Allied Vision for interfacing with their camera modules.

The image capture protocol for the experiments consisted of capturing a 2×2 grid of images that share a 30% overlap centered on the center of each well. For each well, a set of images at multiple vertical planes was also captured. Then, each subunit of a vertical stack was combined with the others in its well using the “Extended Depth of Field (Easy Mode)” plugin in FIJI. These stacks were then stitched together with the others from their grid using the “Pairwise Stitching” plugin with FIJI. These processes were completed in batches using a custom FIJI macro. Image segmentation was performed using the Segment Anything Model (SAM) [23].

### 4.7. Vision-Based Monitoring for Dry run testing

To support the long-term robustness studies described in section 2.3, the system requires a fully automated and repeatable method to verify whether each aspirate or dispense action was executed correctly. Therefore, we developed a vision-based validation pipeline that enables frame-by-frame confirmation of system performance. In each captured frame, two fiducial QR codes printed on the fixture of the system, along with the 24 well regions, are used for geometric registration and alignment. The illumination inside the chamber is uniform and stable, provided by fixed diffused LEDs positioned along the perimeter to minimize surface reflections and ensure consistent imaging conditions.

Once the image is acquired, the processing routine executes locally on the Raspberry Pi. The computational pipeline follows a sequential workflow designed to align, segment, and evaluate each well within the plate. First, the QR codes are detected after a pre-filtering step, and the image is geometrically rectified using ORB (Oriented FAST and Rotated BRIEF) feature matching combined with homography estimation. This alignment step ensures positional consistency across all frames by matching each captured image to a reference template.

After alignment, the frame is cropped to isolate the well region and subsequently subdivided into 24 equal segments corresponding to individual wells. To enhance the visibility of the liquid, image contrast and saturation are adjusted, followed by Gaussian blurring and thresholding in the Lab color space. This sequence isolates the colored medium from the background and generates a binary mask suitable for further analysis.

For each segmented region, a shape similarity metric is computed to estimate how closely the segmented fluid geometry matches the reference geometry. This similarity is evaluated in two steps: first, OpenCV’s matchShapes compares the Hu moments [19] of the current well’s contour against a reference contour, producing a raw dissimilarity score (lower is better); second, an area ratio between the current and reference contour areas is used to penalize size deviations. These two signals are combined into a single normalized similarity score in the range (0, 1], where higher values indicate closer agreement with the reference shape. A tunable threshold is applied to the similarity score to classify each operation as successful or failed.

### 4.8. Cell Culture Conditions

#### mESCs Culture

Mouse embryonic stem cells (mESCs) were dissociated into single cells using TrypLE Express (Thermo Fisher 12604021) for 5 minutes at 37°C. The Mouse Dorsal Organoids were generated as described in [14]. In brief, cells were re-aggregated in Lipidure-coated 96-well V-bottom plates at a density of 3,000 cells per well in 150 L of mESC maintenance medium, supplemented with 10 M ROCK inhibitor Y-27632 (Tocris 1254) and 1,000 U/mL recombinant mouse LIF (Millipore Sigma ESG1107). After 24 h, the medium was replaced with forebrain patterning medium consisting of Dulbecco’s Modified Eagle Medium/Nutrient Mixture F-12 (DMEM/F12) with GlutaMAX (TFS 10565018), 1X chemically defined (CD) lipid concentrate (TFS 11905031), 0.1 mM MEM non-essential amino acids (NEAAs; TFS 11140050), 1 mM sodium pyruvate (MS S8636), 1X N2 supplement (TFS 17502048), and 2X B27 minus vitamin A (-VitA; TFS 12587010). Supplements included 10 M Y27632, 5 M XAV939 (StemCell Technologies (SCT) 1001052), and 5 M SB431542 (Tocris 1614). Medium was changed daily, with N2 and B27 added post-filtration. On day 5, organoids were transferred to ultralow adhesion plates (MS CLS3471) with neuronal differentiation medium and placed on an orbital shaker at 68 rpm. From days 6–12, progenitor expansion medium included a 1:1 mix of Neurobasal-A and BrainPhys media (SCT 05790), supplemented with B27 –VitA, N2 supplement, MEM NEAAs, CD lipid concentrate, and 200 M ascorbic acid (Sigma Aldrich [SA] 49752). Medium was changed every 2–3 days under 5% CO2. From day 13 onward, neural maturation medium consisted of BrainPhys medium supplemented with B27 Serum Free (TFS 17504044), and CD lipid concentrate. Primocin (0.05 mg/mL; InvivoGen antpm05) was included in all media throughout the protocol. For initial mouse experiment, organoids were plated onto the automated cell culture system on day 29, whereas for second experiment, plating was performed on day 18.

#### Immunohistochemistry Analysis

Organoid imaging was performed using a Zeiss AxioImager Z2 microscope equipped with a 10× objective and operated through Zen Blue software. For each sample, z-stack images were acquired at a step size of 1.5 *µ*m across multiple non-adjacent cryosections per organoid.

#### hESCs Culture

Human Embryonic Stem Cells (hESCs) were cultured on 6cm tissue culture-treated plates (Thermo) coated with Vitronectin (Invitrogen) with the stem cell culture media StemFlex (Gibco). Cells were passaged four times using DPBS supplemented with 0.5 mM EDTA (Corning) prior to embryoid body (EB) formation. The H9 human embryonic stem cell line (WA09; WiCell Research Institute; NIH approval # NIHhESC-10-0062) was used under the approval of the UCSF Human Gamete, Embryo, and Stem Cell Research (GESCR) Committee (Protocol 12-08677).

#### Organoid/EB Generation

hESCs are dissociated into single cells using StemPro Accutase dissociation reagent (Gibco). Dissociated cells were centrifuged at 200 RCF to pellet, the supernatant was aspirated, and the pellet was resuspended in AggreWell EB Formation Medium (STEMCELL Technologies) supplemented with 10 *µ*M Y-27632 dihydrochloride, a ROCK inhibitor (Tocris Bioscience). The hESC suspension is quantified using the Invitrogen Countess 3. A total of 3 × 10^6^ cells were plated into a well of the 24-well Aggrewell 800 plate (StemCell Tech), corresponding to approximately 1 × 10^4^ cells per EB. EBs were cultured in AggreWell EB Formation Medium (STEMCELL Technologies) supplemented with SB431542 (Sigma-Aldrich) and IWR1 (EMD Millipore) for 6 days. On day 7, an extracellular matrix (ECM) coating was applied to the EBs. EBs were cultured until day 18 in the same medium. On days 14 and 15, a secondary ECM coating was applied. On day 16, EBs were transferred to the automated cell culture system.

#### Maturation and Medium Change

The AggreWell medium supplemented with SB431542 and IWR1 was further supplemented with penicillin/streptomycin (10,000 U/mL, Gibco) at a 1:100 dilution. The Autoculture system was fed with this medium for 2 days. On day 18, the medium was changed to a Neural Maturation Media (DMEM/F12 + Glutamax, Gibco) + N2 Supplement (Gibco) + penicillin/streptomycin (Gibco) + Amphotericin B (Gibco) + CD Lipid (Gibco).

#### RNA Extraction and qPCR

Organoids were dissociated in 500 *µ*L QIAzol^™^ Lysis Reagent on day 24, and RNA was extracted via ethanol precipitation followed by column purification. cDNA was prepared using the Transcriptor First Strand cDNA Synthesis Kit using 500 ng of RNA. Real-time qPCR was performed on an Applied Biosystems^®^ QuantStudio^™^ 6 Flex Real-Time PCR System using PrimeTime qPCR Master Mix (IDT) with predesigned PrimeTime qPCR probe assays (IDT) targeting FOXG1, PAX6, NKX2.2, TBR1, and B2M. Relative expression was calculated using the 2*^−^*^ΔΔ^*^Ct^* method, normalized to day 30 manually fed control organoids. Osmolarity measurements were performed for all wells in 24-well plate format using a Wescor VAPRO^®^ Model 5600.

## 5. Acknowledgments

This work was supported by the Schmidt Futures Foundation (SF 857; SRS, DH, and MT); the National Human Genome Research Institute (NHGRI) under award RM1HG011543 (SS, DH, and MT); the California Institute for Regenerative Medicine (CIRM) awards DISC4-16285 (SRS, MAM-R and MT) and DISC4-16337 (MAM-R); MZ and YR were funded through the CIRM Fellowship program under award EDUC-12759, the National Science Foundation (NSF) under awards 2134955 (SRS, DH, and MT) and 2034037 (MT); the University of California Office of the President under award M25PR9045 (SRS, MT, and MAM-R); National Institute of Neurological Disorders and Stroke U24NS146314 (MAM-R. and DH); National Science Foundation 2515389 (MT, DH, and MAM-R).

The authors thank the Institute for Stem Cell Biology (IBSC) of UC Santa Cruz and the Life Sciences Microscopy Center (RRID: SCR_021135) for technical support. We also thank Isabel Cline, Samira Vera-Choqqueccota, and John Minnick for their assistance with alternative approaches that were not ultimately used in this work. During the preparation of this work, the authors used AI-based tools to improve the clarity and structure of the sentences. After using this tool, the authors reviewed and edited the content as needed and take full responsibility for the content of the publication.

## Author Contributions

M.Z. conceptualized and engineered the devices and methods, assembled the hardware and control systems, designed and performed experiments, oversaw the project, and wrote the manuscript. D.E. and Y.R. developed the internet-connected microscopy system. M.P.M. developed the software infrastructure, user interface, computer vision analyses, and image-based validation methods. S.H., H.E.S., and D.O. prepared biological samples and performed biological experiments, analysis, and data interpretation. E.W. contributed to PCB design. S.T.S. and K.V. helped develop the software and set up the syringe pump. C.J., M.A.M.-R., S.R.S., and D.H. provided guidance on experimental design, data interpretation, and analysis. M.T. conceptualized, supervised the work, and contributed to the experimental methodology, data interpretation, and manuscript writing.

## Supplementary Materials

Supplementary figures: Figures S1–S6

Supplementary tables: Tables S1–S2

Supplementary text related to Figures S2 and S5

Supplementary videos: Videos S1–S3

## Data availability

All data supporting the findings of this study are available within the paper and its Supplementary Information.

## Code availability

The user interface and control software developed for experiment scheduling, device control, real-time monitoring, and data acquisition are openly available under a CC-BY-4.0 license on Zenodo at https://doi.org/10.5281/zenodo.20673645 (v1.0.0) [27].

