## Supplemental Materials for "Servo-Actuated 3D-Printed Disposable Microvalves for Automated, Scalable Organoid Culture in Standard Incubators"

---

---

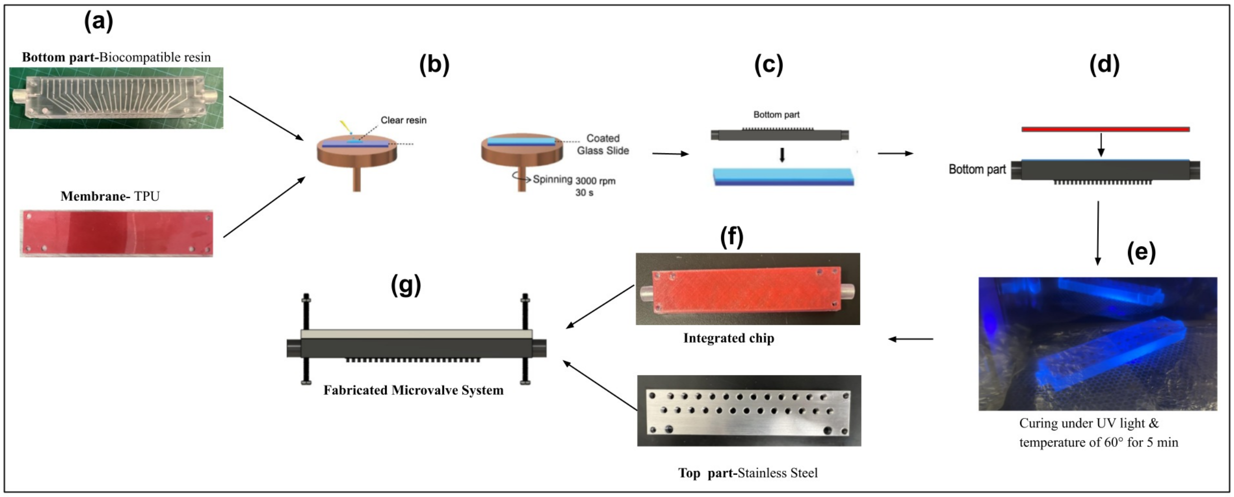


**Figure S1:** Fabrication process of the microfluidic microvalve. **(a)** SLA 3D-printed bottom component fabricated using a transparent biocompatible resin (BioMed clear resin) and an FFF 3D-printed TPU membrane. **(b)** Formation of a thin resin layer (BioMed clear resin) on a glass substrate using spin coating. **(c)** Placement of the bottom component onto the resin-coated glass surface. **(d)** Stamping of the TPU membrane onto the bottom component with a thin resin film. **(e)** UV curing of the assembled components at 60 °C for 5 min. **(f)** Fabricated integrated microfluidic component together with the top component manufactured from stainless steel using CNC milling. **(g)** Assembled microvalve system.

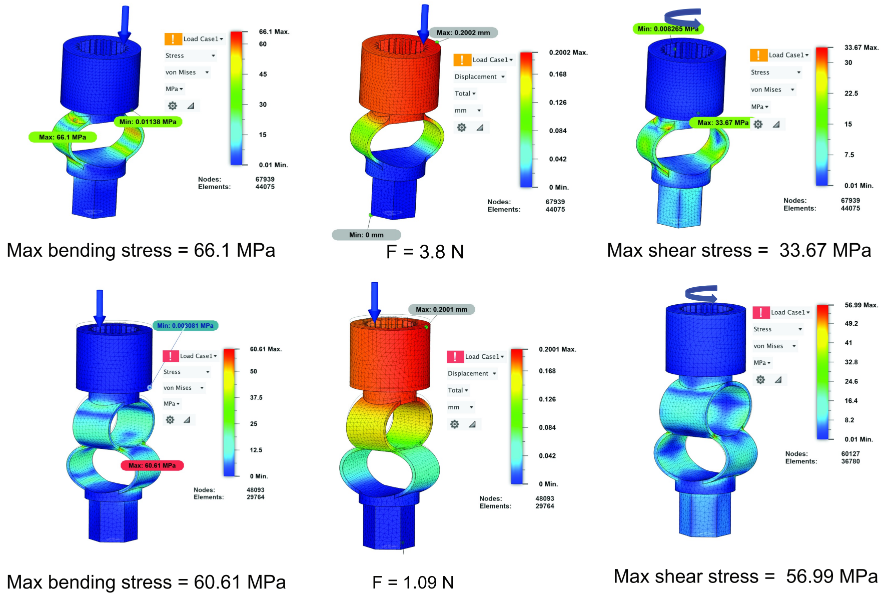


**Figure S2:** Finite element analysis of candidate adapter designs for coupling the servo shaft to an M5 screw under applied torque. Bending stress, force, and shear stress distributions were evaluated to identify geometries that minimize loading on the servo shaft. The optimized designs combine flexibility in the Z-direction and rigidity in the XY-plane.

Finite element analysis (FEA) was performed to evaluate ten design candidates for coupling the servo shaft to an M5 screw. Simulations were conducted under the maximum servo torque (7.3 N·mm) and the required displacement for membrane actuation (200 µm). The objective was to identify geometries that minimize mechanical loading on the shaft while maintaining sufficient rigidity for effective torque transmission.

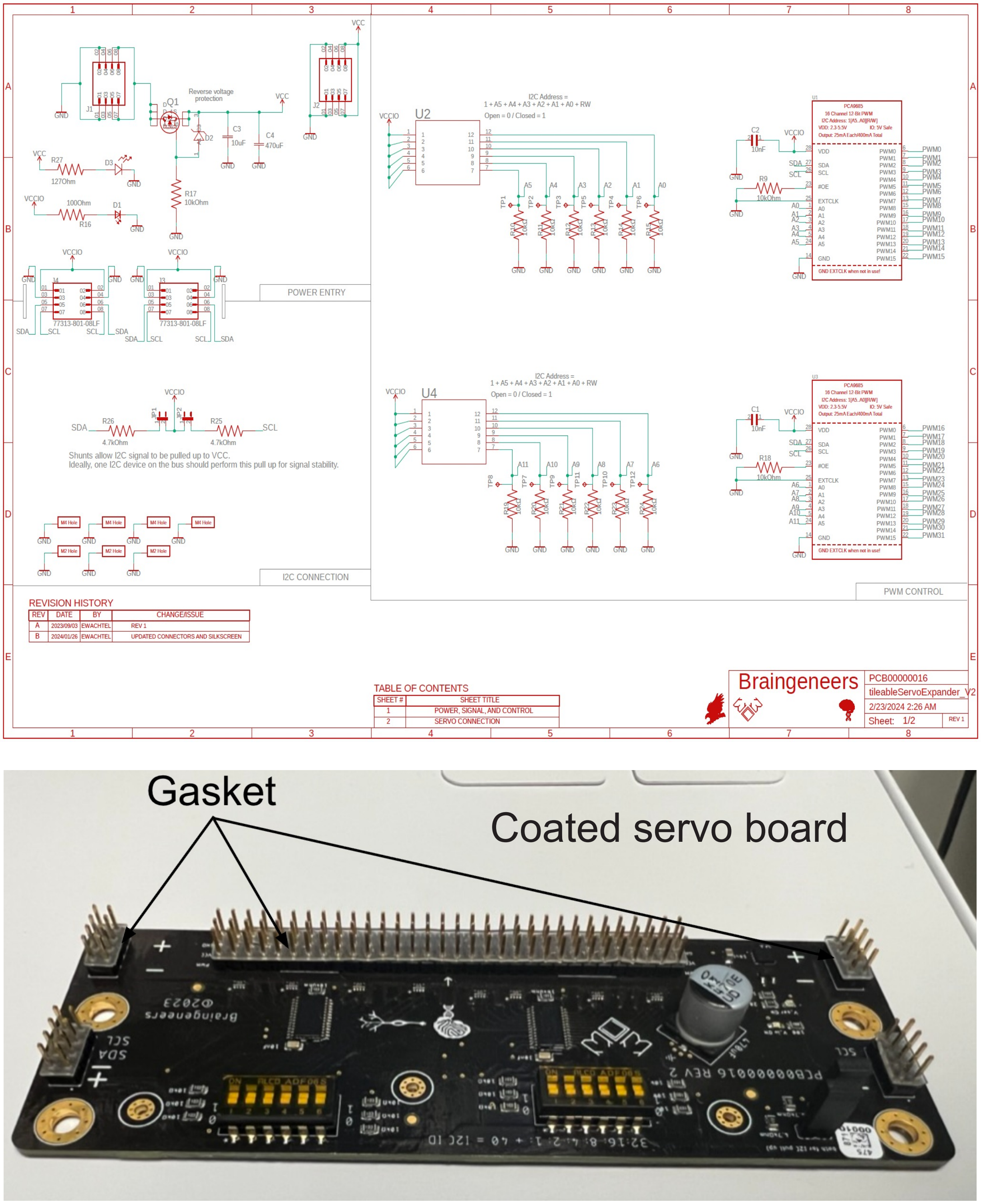


**Figure S3:** Custom control boards used in the automated cell culture system. Schematic diagram of the servo control board. Custom-designed servo control board capable of supporting up to 32 servos; the board is coated with Super Corona Dope (MG Chemicals) and silicon gaskets are used for the pins to ensure electrical insulation under high-humidity conditions.

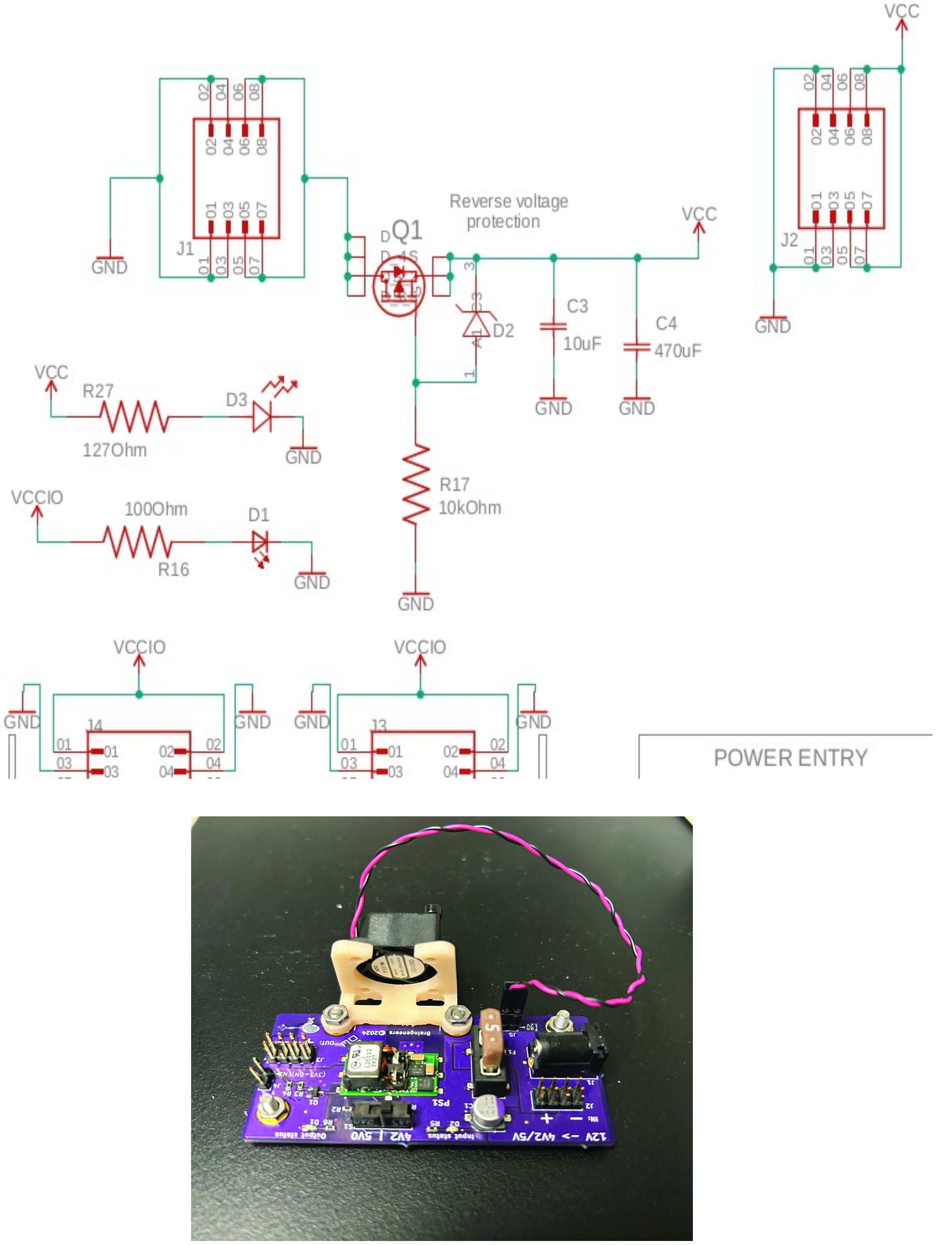


**Figure S4:** Custom control boards used in the automated cell culture system. Power switching (power-cut) circuit for controlling servo supply.

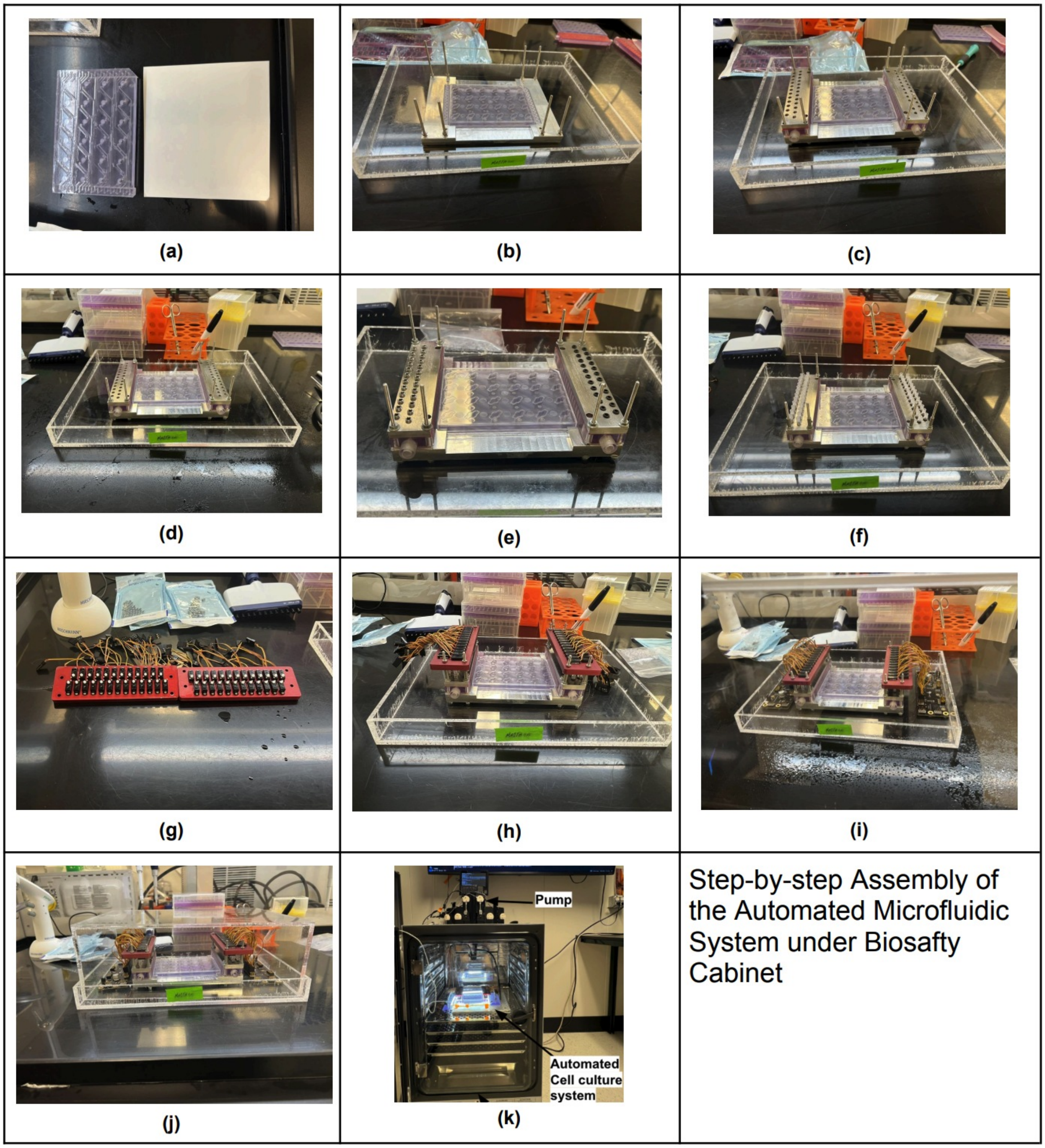


**Figure S5:** Assembly process of the automated microfluidic system under sterile conditions

**(a) Apply adhesive tape:** Cut a piece of microfluidic adhesive tape (9795R, 3M) to match the size of the 24-well plate and apply it to the bottom surface. Ensure that no air bubbles are trapped between the tape and the printed chip.

**(b) Insert the well plate:** Place the 24-well plate into the main metallic fixture (base part) of the system. Insert long threaded rods (M3 × 0.5 × 80 mm, 304 stainless steel) into each corner of the fixture. Place two laser-cut gaskets between the well plate and the microvalve assembly to ensure proper sealing of the ports.

**(c) Install dispensing and aspiration valves:** Assemble the fluidic components with the stainless steel parts containing threaded holes and secure them using four M3 threaded rods.

**(d) Secure valves to fixture:** Fasten the dispensing and aspiration valve assemblies to the metallic fixture using four M3 nuts (94777A105, M3 $\times$ 0.5 mm, McMaster-Carr) on each side. Use two additional M3 screws (93190A125, McMaster-Carr) and corresponding M3 nuts to ensure leak-free sealing between the plate and valve interfaces.

**(e) Insert M5 ball-tip screws:** Insert 25 M5 ball-tip screws (M5-0.8 $\times$ 8 mm, NACX) on each side of the system.

**(f) Add spacers and M5 adapters:** Placing the 3D-printed M5 adapters and spacers (92871A304, McMaster-Carr).

**(g) Attach servomotors:** Mount the 50 servomotors onto their holders using M1.6 screws (99461A911, McMaster-Carr). The servo holders are CNC-machined from AL6061 and anodized (Billet Metal Craft, Santa Cruz, CA, USA).

**(h) Install servo holder assembly:** Install the servo holder assembly as shown in Fig. (h), ensuring proper alignment of the coupling mechanism with the servo shafts. Secure the assembly to the main fixture using four M3 nuts (94777A105, M3 $\times$ 0.5 mm, McMaster-Carr).

**(i) Connect to circuit boards:** Connect each servomotor to the corresponding circuit board pins.

**(j) Transfer to incubator:** Place the fully assembled system in a sterile container (j) to transfer it from the biosafety cabinet to the incubator, and connect it to the syringe pumps (k).

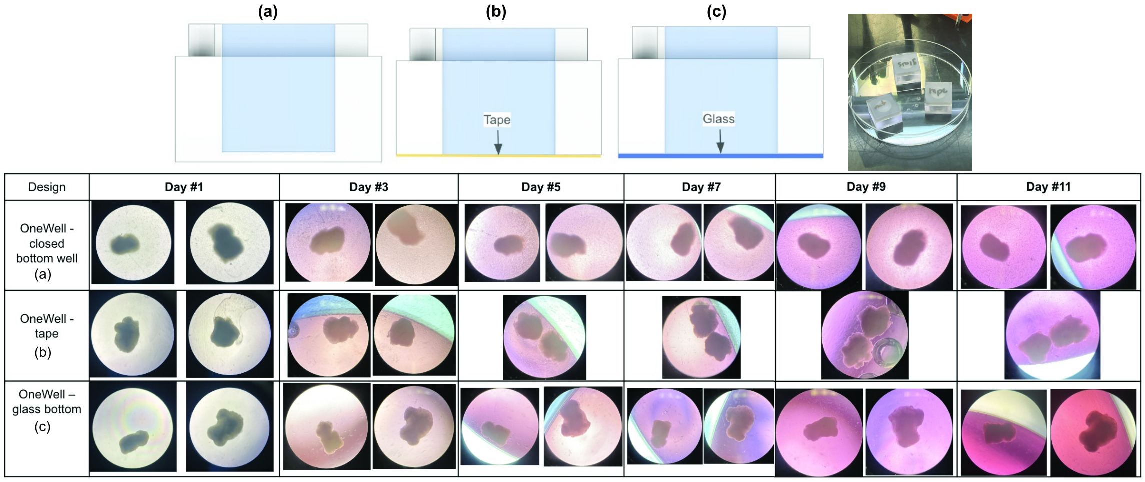


**Figure S6:** Biocompatibility assessment of Detax FREEPRINT Ortho resin for organoid culture

**Table S1:** Dry-run validation of automated media handling performance across wells, assessing reliability and consistency of aspiration and dispensing.

| Well# | Aspiration Commands | Successful Aspiration | Failed Aspiration | Dispensing Commands | Successful Dispensing | Failed Dispensing | Success rate Aspiration% | Success rate Dispensing% |
|---|---|---|---|---|---|---|---|---|
| **A1** | 97 | 97 | 0 | 97 | 97 | 0 | 100.00 | 100.0 |
| **A3** | 97 | 97 | 0 | 97 | 97 | 0 | 100.00 | 100.0 |
| **A4** | 97 | 95 | 2 | 97 | 96 | 1 | 97.94 | 99.0 |
| **A6** | 97 | 97 | 0 | 97 | 97 | 0 | 100.00 | 100.0 |
| **B1** | 96 | 96 | 0 | 96 | 96 | 0 | 100.00 | 100.0 |
| **B2** | 97 | 96 | 1 | 97 | 96 | 1 | 98.97 | 99.0 |
| **B3** | 97 | 95 | 2 | 97 | 97 | 0 | 97.94 | 100.0 |
| **B4** | 95 | 95 | 1 | 95 | 94 | 1 | 100.00 | 98.9 |
| **B5** | 95 | 94 | 1 | 95 | 95 | 1 | 98.95 | 100.0 |
| **B6** | 96 | 96 | 0 | 96 | 96 | 0 | 100.00 | 100.0 |
| **C1** | 96 | 96 | 0 | 96 | 96 | 0 | 100.00 | 100.0 |
| **C2** | 95 | 95 | 0 | 95 | 95 | 0 | 100.00 | 100.0 |
| **C3** | 96 | 96 | 0 | 96 | 96 | 0 | 100.00 | 100.0 |
| **C4** | 92 | 84 | 8* | 92 | 92 | 0 | 91.30 | 100.0 |
| **C5** | 85 | 78 | 9* | 85 | 85 | 0 | 91.76 | 100.0 |
| **C6** | 97 | 94 | 3 | 97 | 97 | 0 | 96.91 | 100.0 |
| **D1** | 96 | 95 | 1 | 96 | 96 | 0 | 98.96 | 100.0 |
| **D2** | 97 | 97 | 0 | 97 | 97 | 0 | 100.00 | 100.0 |
| **D3** | 96 | 88 | 8* | 96 | 96 | 0 | 91.67 | 100.0 |
| **D4** | 96 | 96 | 0 | 96 | 96 | 0 | 100.00 | 100.0 |
| **D5** | 96 | 90 | 6* | 96 | 96 | 0 | 93.75 | 100.0 |
| **D6** | 96 | 95 | 1 | 96 | 96 | 1 | 98.96 | 100.0 |

* Aspiration failures occurred due to temporary sticking of the membrane to the valve seat. A brief positive pressure step applying prior to aspiration helping with membrane release.

**Table S2:** Osmolarity measurements

| Well | Medium | Feeding Protocol | Osmolarity (mmol/kg) |
|---|---|---|---|
| **Sample** | DMEM/F12 + Glutamax + N2 | Fresh | 290 |
| **A2** | DMEM/F12 + Glutamax + N2 | LFO1* | 410 |
| **A3** | DMEM/F12 + Glutamax + N2 | LFO1* | 437 |
| **A4** | DMEM/F12 + Glutamax + N2 | LFO2* | 342 |
| **A5** | DMEM/F12 + Glutamax + N2 | LFO2* | 374 |
| **A6** | DMEM/F12 + Glutamax + N2 | LFO2* | 335 |
| **B2** | DMEM/F12 + Glutamax + N2 | LFI1* | 332 |
| **B3** | DMEM/F12 + Glutamax + N2 | LFI1* | 411 |
| **B4** | DMEM/F12 + Glutamax + N2 | LFI2* | 338 |
| **B5** | DMEM/F12 + Glutamax + N2 | LFI2* | 331 |
| **B6** | DMEM/F12 + Glutamax + N2 | LFI2* | 341 |
| **C1** | DMEM/F12 + Glutamax + N2 | HFI1* | 329 |
| **C2** | DMEM/F12 + Glutamax + N2 | HFI1* | 328 |
| **C3** | DMEM/F12 + Glutamax + N2 | HFI1* | 325 |
| **C4** | DMEM/F12 + Glutamax + N2 | HFI2* | 326 |
| **C5** | DMEM/F12 + Glutamax + N2 | HFI2* | 329 |
| **C6** | DMEM/F12 + Glutamax + N2 | HFI2* | 327 |
| **D1** | DMEM/F12 + Glutamax + N2 | HFO1* | 355 |
| **D2** | DMEM/F12 + Glutamax + N2 | HFO1* | 349 |
| **D3** | DMEM/F12 + Glutamax + N2 | HFO1* | 339 |
| **D4** | DMEM/F12 + Glutamax + N2 | HFO2* | 342 |
| **D5** | DMEM/F12 + Glutamax + N2 | HFO2* | 337 |
| **D6** | DMEM/F12 + Glutamax + N2 | HFO2* | 339 |

* LFO = Low Frequency Outer row (A), LFI = Low Frequency Inner row (B), HFI = High Frequency Inner row (C), and HFO = High Frequency Outer row (D).
